# Microbiological water quality of Managed Aquifer Recharge systems in the salinity-prone southwest coastal Bangladesh

**DOI:** 10.1101/2020.03.02.972372

**Authors:** Solaiman Doza, Abu Mohd Naser, Md Mahbubur Rahman, Momenul Haque Mondol, Golam Kibria Khan, Md. Nasir Uddin, Mohammed Shahid Gazi, Gazi Raisul Alam, Mohammed Rabiul Karim, Kazi Matin Ahmed, Stephen P. Luby, Thomas Clasen, Leanne Unicomb

## Abstract

Managed aquifer recharge (MAR), a hydro-geological intervention designed to dilute groundwater salinity, pumps pond water treated through a slow sand filter into the underground aquifers. We evaluated the microbiological safety of the resulting MAR water at sites from three districts in southwest coastal Bangladesh. We collected monthly paired pond-MAR water samples from July 2016-June 2017 and enumerated fecal coliforms and *E. coli* using the IDEXX quanti-tray technique, by the most probable number (MPN) method. We used WHO risk categories for microbiological quality; no risk (<1 MPN), low risk (1-10 MPN) and moderate to high risk (>10 MPN per 100 mL water). We estimated the difference in mean log_10_ MPN in pond and MAR water using linear mixed effect models with random intercepts and cluster adjusted robust standard error. Almost all pond water samples (292/299, 98%) had moderate- to high-risk level (>10 MPN) fecal coliforms and *E. coli* (283/299, 95%). In contrast, 81% (242/300) of MAR water samples had no or low risk level fecal coliforms (0-10 MPN), of which 60% (179/300) had no fecal coliforms. We detected no or low risk level *E. coli* in 94% (283/300) of MAR water samples of these 80% (240/300) had no *E. coli.* MAR samples had lower mean log_10_ MPN fecal coliforms (-2.37; 95% CI: -2.56, -2.19) and *E. coli* (-2.26; 95% CI: - 2.43, -2.09) than pond water; microbial reductions remained consistent during the wet (May-October) and dry seasons. MAR-systems provided water with reduced fecal indicator bacteria compared to infiltered pond water.

## Background

Reliance on unsafe drinking water is associated with infection and disease, including childhood diarrhea, a leading causes of morbidity and mortality in low and middle income countries ^1-4^. A recent study in rural Bangladesh reported increased prevalence of diarrhea (prevalence ratio = 1.14, 95% CI = 1.05, 1.23) associated with each 10-fold increase in *E. coli* count in their drinking water ^5^. To reduce the risk of waterborne disease rainwater harvesting and groundwater has been widely promoted over surface water ^6, 7^. The aquifers in the southwest coastal districts of Khulna, Bagerhat and Satkhira are frequently saline, providing brackish water that greatly hinders freshwater access for the surrounding communities ^8-10^. While many householders practice rainwater harvesting during the monsoon months of May through October, limited storage capacity requires them to revert to pond water during dry seasons ^8^.

Palatable drinking water was lacking especially during the dry months in the salinity affected southwest coastal communities. Researchers from Geology Department of the University of Dhaka designed managed aquifer recharge (MAR), an intervention to artificially dilute the groundwater salinity by pumping, treating and infiltrating pond water into the aquifer ^11,12^. The recharged water remains protected from evaporation since freshwater is infiltrated via wells beneath layers of clay ^12^.In collaboration with UNICEF, they piloted 20 MAR projects between 2009-2012. The MAR systems were constructed with local resources and managed by trained locals. The sites had an average storage of approximately 900 m^3^ of fresh water per year, sufficient to deliver 15 L of safe drinking water/day to community members ^13^. The pilot research showed that the salinity of water drawn from surrounding hand pumps was reduced to acceptable limits (≤2 mS/cm) in 16 of the 20 MAR sites ^11, 14^. Based on the successful pilot of 20 MAR systems a larger project was initiated, and 75 new MAR systems were installed between 2013 and 2014 in three coastal districts (Satkhira, Khulna and Bagerhat) ^14^. To evaluate the health effects of MAR water we conducted a stepped wedge cluster-randomized controlled trial ^15^.

Groundwater is generally free of microbiological pathogens as inactivation and reduction occurs naturally while water percolates through the soil layers ^16^. Nevertheless, recent studies in Bangladesh reported that up to 65% of tubewells with handpumps contain low levels of fecal contamination ^17-19^. Tubewells located near latrines, lakes or ponds can be fecally contaminated by unsealed handpumps or leaks at the base ^20^. MAR systems use a shallow well with handpump for water abstraction, creating a risk of fecal contamination through surface runoff from unprotected wells ^21^. The MAR systems utilized slow sand filters to reduce the risk of fecal pathogen transmission that are common in surface water. During piloting, MAR systems successfully reduced *E. coli* in the abstracted water though half of the samples still had low *E.coli* counts which might be introduced from the pond water or the surface runoffs ^15, 21, 22^. In the present study, our aim was to evaluate the new MAR systems’ microbiological safety throughout the year. We collected and tested both MAR and source pond waters for fecal coliforms and *E.coli* once/per month for 12 months.

## Materials and Methods

### MAR systems

During MAR system piloting, several designs were considered; the design found both cost-effective and efficient for water storage was selected for scaling up ^14, 21^. Ponds were the freshwater sources, pumped into an overhead concrete reservoir comprised two chambers, the first one for filtration and the second for storage. The water inlet at the pond had a screen to prevent fish or unwanted debris entering the filtration chamber. The pumped water percolated through a slow sand filter, passed into infiltration wells by gravitational force; there was no chemical treatment of the infiltration water. Four infiltration wells were placed in a square to create a fresh water bubble at the top of the brackish aquifer ^21^. The abstraction well was placed in the middle of the infiltration wells and was connected to a hand pump from which people collected drinking/cooking water. No latrine was present within 10 meters of the MAR systems.

### Study sites

The UNICEF-University of Dhaka team constructed 75 new MAR sites. Sites were selected based on groundwater salinity, measured as electrical conductivity (EC), was ≤2 mS/cm ^2, 3, 14^. We purposefully selected the first 25 sites that met the EC level criterion. These were located in the sub-districts of three coastal districts; 9 sites in Satkhira, 7 sites in Khulna, and 9 sites in Bagerhat (Figure 1). MAR site and aquifer characteristic data were provided by the UNICEF-University of Dhaka team.

**Figure 1:**
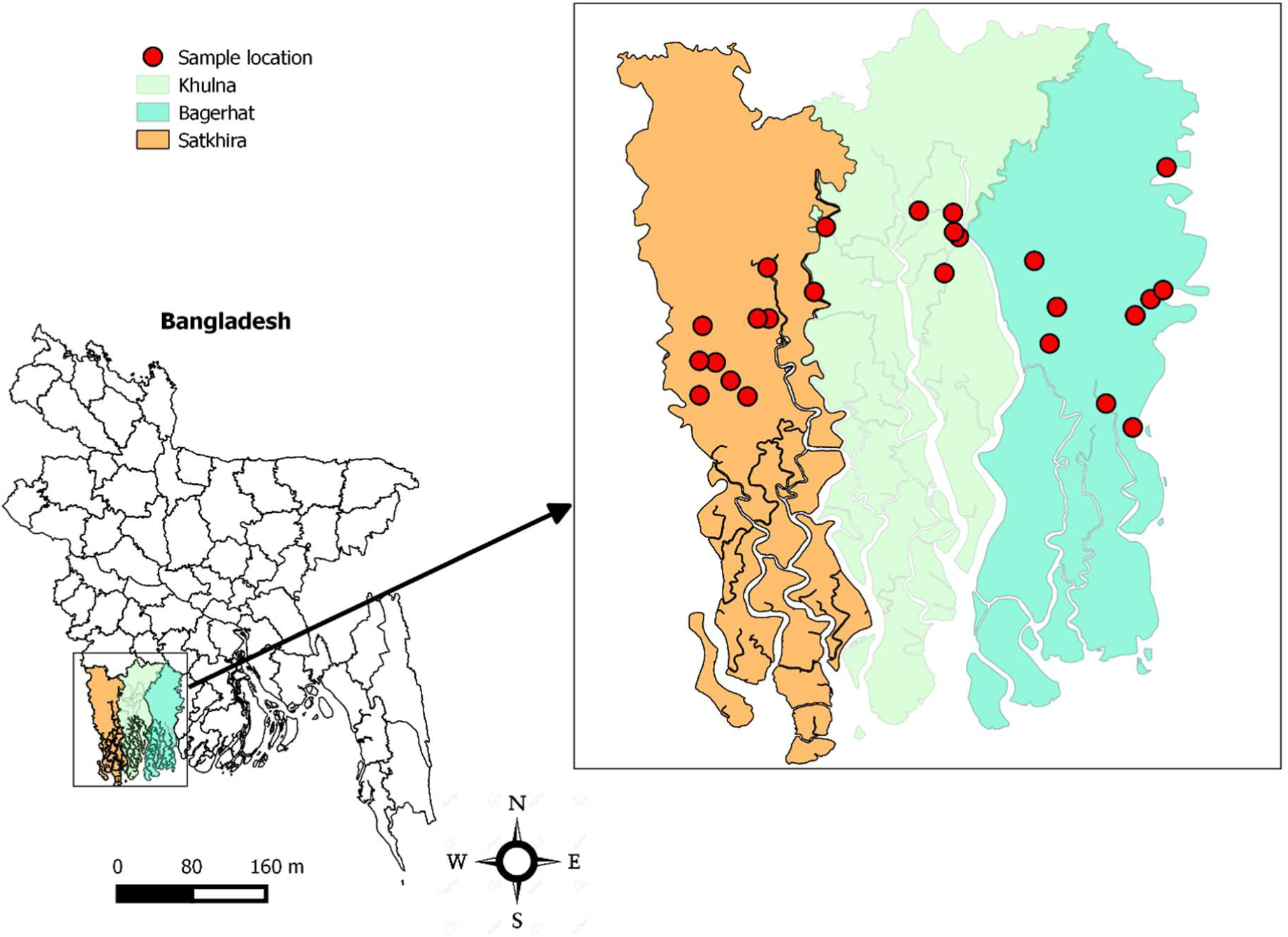
Selected MAR sites for the current study located in coastal districts of Bangladesh.

### Water sample collection and microbial testing

Before initiating sample collection, field and laboratory assistants were trained following standard operating procedures for sample collection and testing. From July 2016, field assistants collected 100 mL samples from both the source ponds and the abstraction wells. To collect pond water, they switched on the water pump that moved pond water into the overhead reservoir and collected the sample from the inlet that drained into the filter chamber. For MAR water collection, they purged the community hand-pump to discard stagnant water (20-30 L) from the well pipe. Samples were collected aseptically after the tubewell head was disinfected by wiping with 70% ethanol solution. All samples were collected in a sterile Whirlpak bag^®^(Nasco Modesto, Salida, CA) and placed in a cooler box maintaining 4-8 °C. The field assistants transported samples to the field laboratory in Khulna within 6 hours of collection.

In the lab, we made 1:1 dilutions for both pond and MAR water (50 mL of the raw water and 50 mL distilled water mixed in the dilution bag) and tested the sample using the IDEXX quanti-tray technique with Colilert-18 media (IDEXX Laboratories, Inc.) for enumeration of fecal coliforms and *E. coli*. Sample trays were incubated at 44.5 °C for 18-22 hours. Both fecal coliforms and *E. coli* were enumerated using the most probable number method (MPN) following the manufacturer’s instruction. We ran laboratory blanks on each day of sample testing and field blanks (a sample bag filled at the field site with sterile distilled water) once a week to monitor cross-contamination during collection or processing. On every third day of sample collection, a duplicate sample was collected and tested and once each week laboratory replicates (two aliquots from the same sample) were included to ensure sample quality. None of the laboratory blanks were positive for fecal coliforms and only one of the field blanks was positive for fecal coliforms (MPN count was 3.1).

### Data analysis

We estimated the proportion of pond and MAR samples that had fecal coliforms and *E. coli* per 100 mL water. We replaced the zero-count data with 0.5 MPN/100ml and log_10_ transformed the MPN count to minimize the skew in distribution. To analyze seasonal variation, we categorized the year into dry and wet months; rain generally starts from May and lasts until October which was defined as wet season and November to April was categorized as the dry season ^23^. We estimated the mean log_10_ MPN of fecal coliforms and *E. coli* and their mean log_10_ differences between pond and MAR water by using the linear mixed effect model with random intercept to adjust for clustering effects of 12-month follow-ups and MAR-pond pairs. We also adjusted the estimated effects for seasonal variation. We analyzed contamination in three categories based on WHO risk categories: no risk (<1 MPN/100 mL), low risk (1 to <10 MPN/100 mL) and moderate to high risk (>10 MPN/100 mL) ^2, 23^. To compare the proportions of samples with each water quality category, we used the cumulative of ordered logistic regression with multilevel mixed effect model ^24^. To explore factors associated with moderate- to high-risk MAR water contamination, we used conditional logistic regression for adjusting pairs (MAR water and input pond water from each MAR site), and examined aquifer characteristics, season and pond fecal coliforms and *E. coli* contamination. We performed correlation analysis for MAR vs. pond fecal coliforms and *E. coli* MPN/100ml. We performed all statistical analysis in STATA (version 13.0). We plotted the mean log_10_ MPN fecal coliforms and *E. coli* over month using the line diagrams with standard errors in R software (version 3.4.3).

## Results

### Location and geological variations across the study sites

Similar numbers of sites were selected from the three study districts (9 from Bagerhat, 7 from Khulna and 9 from Satkhira) and there was little variation in terms of aquifer size; in Bagerhat, the median aquifer thickness was 12m (IQR: 12-14) in contrast to 14m (IQR: 11-15) in Khulna and 11m (IQR: 9-15) in Satkhira. The median aquifer depth from the surface was slightly lower in Bagerhat (18m; IQR: 17-24) than Khulna (23m; IQR: 17-25) and Satkhira (21m; IQR: 17-25). Clay layer thickness or the depth from the surface plays a key role for natural infiltration of surface runoffs and facilitates rapid microbial transmission. In Satkhira, the median clay thickness was the highest (18m; IQR: 11-24). Bagerhat and Khulna sites had similar top clay layers with median thickness of 11m (IQR: 8-15) and 12m (IQR: 9-20) respectively (Table 1).

**Table 1:** Geological variations in 25 MAR sites across study districts.

|  | Districts |  |  |
| --- | --- | --- | --- |
|  | Bagerhat | Khulna | Satkhira |
| Site % (n) | 36 (9) | 28 (7) | 36 (9) |
| Aquifer thickness/size (m), median (IQR) | 12.2 (12.2-13.7) | 13.7 (10.7-15.2) | 10.7 (9.1-15.2) |
| Aquifer depth from surface (m), median (IQR) | 18.3 (16.8-24.4) | 22.9 (16.8-27.4) | 21.3 (16.8-27.4) |
| Clay thickness from surface (m), median (IQR) | 10.7 (7.6-15.2) | 12.2 (9.1-19.8) | 18.3 (10.7-24.4) |

### Fecal contamination in MAR water compared to source water

The year-round mean log_10_ MPN of fecal coliforms was 0.28 (95% Confidence Interval (CI): 0.12, 0.43) in the MAR water versus 2.65 (95% CI: 2.50, 2.81) in pond water, a difference of -2.37 (95% CI: -2.56, -2.19) (Table 2). The year-round mean log_10_ MPN of *E. coli* was -0.10 (95% CI: -0.20, 0.08) in MAR water versus 2.20 (95% CI: 2.10, 2.34 in pond water, a difference of -2.26 (95% CI: -2.43, -2.09). Almost all pond water samples had levels of fecal coliforms (292/299, 97.7%) and *E. coli* (283/299, 94.7%) in excess of 10 MPN, rendering the moderate to high risk. In contrast, 179 (59.7%) of the 300 MAR well samples had no risk (<1 MPN) and 58 (19.3%) had moderate to high risk. We detected 80% (240/300) of MAR water had no risk (<1 MPN) for *E. coli*, and 17 (5.7%) had moderate to high risk counts (>10 MPN). The log-odds of moderate to high risk counts compare to combined low and no risk counts was 5.42 (95% CI: 4.4, 6.5; p<0.01) for the fecal coliforms and 5.87 (95% CI: 4.9, 6.9; p<0.01) for *E. coli* in MAR water samples than the pond water (Table 3).

**Table 2:**
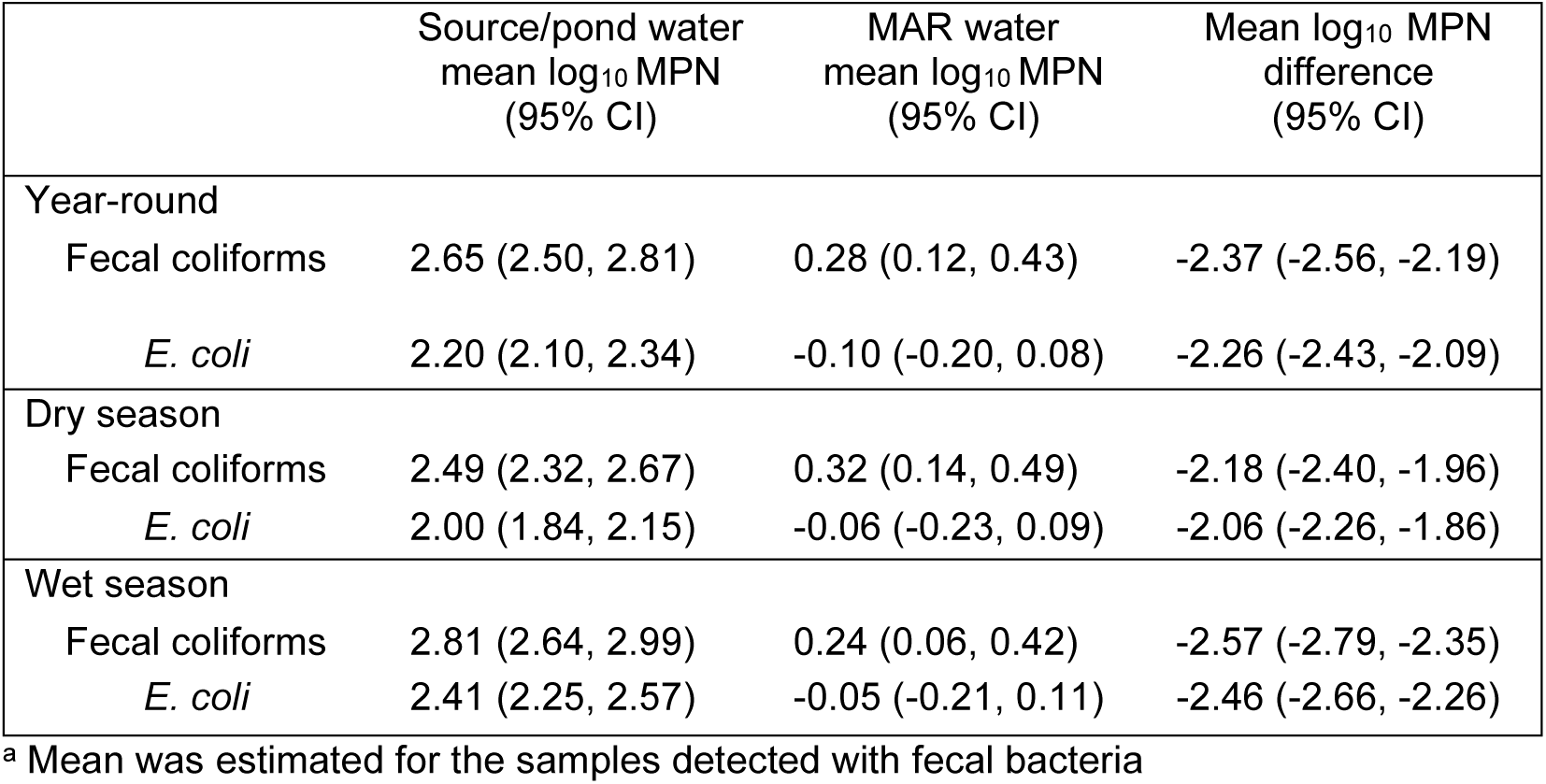
Mean log_10_ MPN reduction in MAR water.

|  | Source/pond water<br>mean log<sub>10</sub> MPN<br>(95% CI) | MAR water<br>mean log<sub>10</sub> MPN<br>(95% CI) | Mean log<sub>10</sub> MPN<br>difference<br>(95% CI) |
| --- | --- | --- | --- |
| Year-round |  |  |  |
| Fecal coliforms | 2.65 (2.50, 2.81) | 0.28 (0.12, 0.43) | -2.37 (-2.56, -2.19) |
| *E. coli* | 2.20 (2.10, 2.34) | -0.10 (-0.20, 0.08) | -2.26 (-2.43, -2.09) |
| Dry season |  |  |  |
| Fecal coliforms | 2.49 (2.32, 2.67) | 0.32 (0.14, 0.49) | -2.18 (-2.40, -1.96) |
| *E. coli* | 2.00 (1.84, 2.15) | -0.06 (-0.23, 0.09) | -2.06 (-2.26, -1.86) |
| Wet season |  |  |  |
| Fecal coliforms | 2.81 (2.64, 2.99) | 0.24 (0.06, 0.42) | -2.57 (-2.79, -2.35) |
| *E. coli* | 2.41 (2.25, 2.57) | -0.05 (-0.21, 0.11) | -2.46 (-2.66, -2.26) |

<sup>a</sup> Mean was estimated for the samples detected with fecal bacteria

**Table 3:** Fecal contamination in MAR and source water, 2016-2017.

| Indicator<br>bacteria<br>(WHO<br>disease risk<br>category)* | Source water<br>N = 299 |  |  | MAR water<br>N = 300 |  |  | Difference between source<br>and MAR water |  |
| --- | --- | --- | --- | --- | --- | --- | --- | --- |
|  | <1 MPN<br>(no risk) | 1-10<br>MPN<br>(low risk) | >10 MPN<br>(moderate<br>to high risk) | <1<br>MPN<br>(no<br>risk) | 1-10<br>MPN<br>(low risk) | >10 MPN<br>(moderate<br>to high risk) | Change in<br>ordered log-odds of higher<br>risk category<br>(OR) | 95% CI |
| Fecal<br>coliforms | 3 (1) | 4 (1.3) | 292 (97.7) | 179<br>(59.7) | 63 (21.0) | 58 (19.3) | -5.42<br>(0.004) | -6.45, -4.40<br>(0.002,<br>0.012) |
| *E. coli* | 8 (2.7) | 8 (2.7) | 283 (94.7) | 240<br>(80.0) | 43 (14.3) | 17 (5.7) | -5.87<br>(0.003) | -6.87, -4.88<br>(0.001,<br>0.008) |

\* WHO guideline for drinking water <sup>2</sup>;

The mean log_10_ MPN *E. coli* difference between MAR and pond water was significantly greater (p<0.05) during the wet season (−2.46, 95% CI -2.66, -2.26) compared to dry (−2.06, 95% CI -2.26, -1.86). For mean log_10_ MPN fecal coliform concentrations, the difference was similarly higher (wet season: -2.57, 95% CI -2.79, -2.35 versus dry season: -2.18, 95% CI -2.40, -1.96) but was not significant (Table 2). The mean log_10_ MPN fecal coliform concentrations in MAR water was consistently lower throughout the study months with a slight increase in July-16, Nov-16, Jan-17, and Apr-Jun-17. The mean log_10_ MPN *E. coli* counts in MAR water were slightly higher in July-16, decreased in August-16 and remained similar throughout the study period (Figure 2).

**Figure 2:**
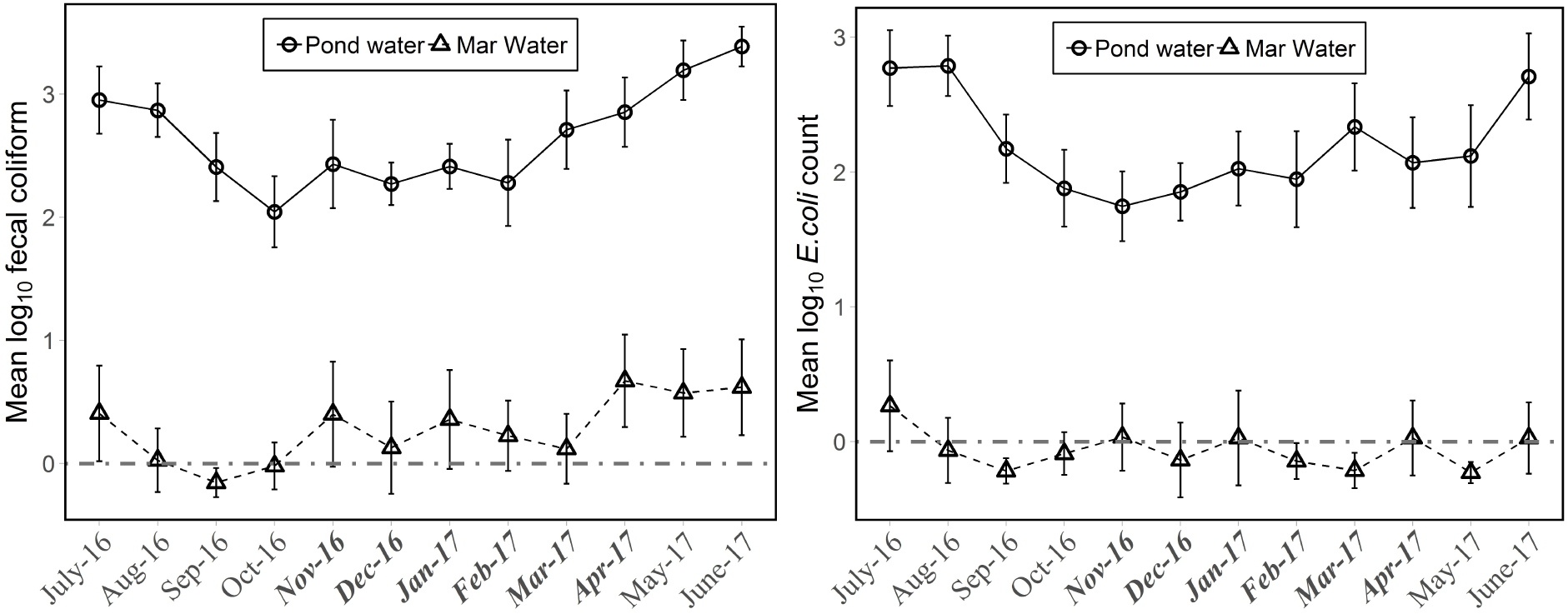
Mean log_10_ MPN fecal coliform and *E. coli* difference between source/pond and MAR water (Note: bold italic font indicates dry season month)

Of the 10 sample-pairs, 8 were collected during the dry season. There were 3 sites for which there was more than one month where the MAR sample had higher fecal coliform levels than pond water. For one site this occurred during three consecutive months (January to March 2017), during two of which *E. coli* contamination levels were similarly higher for MAR water than pond water (January, February 2017). For *E. coli*, there were 6 paired pond-MAR water samples where counts were higher in MAR water than pond water samples, excluding those where both samples had no *E. coli*. Among these 6, two came from the same site and consecutive months (data not shown). There was a significant positive correlation between MAR and pond water log_10_ MPN fecal coliforms (r=0.24, p=0.000) and a positive borderline significant correlation for *E. coli* (r=0.1, p=0.054).

## Discussion

The MAR systems in the southwest coastal Bangladesh delivered water that mostly had no detectable fecal contamination throughout a year. Both fecal coliforms and *E. coli* were significantly reduced to no or low risk contamination levels.

Fecal contamination is commonplace during the monsoon particularly due to increased water levels and saturated soil conditions which permits microbes to move freely and travel longer distances ^25, 26^. Similarly, MAR source ponds had increased fecal load during the wet months, but the MAR systems successfully retained water with no to low risk fecal contamination. Shallow aquifers (<50 meters) often have low-level fecal contamination due to proximity to contamination sources such as pit latrines or intrusion from highly contaminated surface water bodies ^27^. But the MAR systems were constructed at a safe distance from latrines and in areas with confined aquifers that had a top clay layer ^14, 21^. Clay layers generally hinder natural infiltrations and thus preserve the brackish water throughout the year. We anticipate this also protected MAR waters from fecal contamination especially during the wet months when environmental fecal load increases as do source ponds ^19^.

The MAR systems used a slow sand filter to remove microorganisms from the source water prior recharging into the underground aquifer, storing water beneath the ground which may reduce opportunities for pathogen die-off through natural process. Slow sand filters are well-known for their effectiveness in reducing fecal bacteria, yet they also have certain limitations. For example, many pond sand filters located in rural southwest coastal Bangladesh became clogged or broke down due to irregular maintenance and became abandoned. Lack of community participation was an important contributor to the failure of pond sand filters ^14, 28^. MAR systems had similar management challenges such as the risk of a broken pump or inadequate power for pumps that needed fuel or electricity to function. The appointed caretakers from the respective communities were responsible for keeping the MAR systems operational. We did not collect the MAR maintenance data and thus unable to predict the site management scenario. However, the consistency in microbiological quality of MAR water suggests the overall maintenance was adequate to deliver optimum quality drinking water throughout the year.

The study sampled from 25 of the 75 sites therefore we cannot assume that other sites had similar quality water and may not represent MAR systems with adverse hydrogeological conditions. However, we collected samples for 12 months from all three districts, and these mostly represented geological and geographical variations in the study region. We did not explore the maintenance of the MAR systems sites and this may have had an impact on MAR system performance. A major drawback of community led water treatment systems in the past has been poor maintenance induced by lack of ownership. In Bangladesh, pond sand filters were successful for initial periods but broke down later and left behind by the communities ^14^. MAR systems require regular freshwater recharge using electricity or fuel to pump water, the sand filter needs monthly cleaning and maintenance thus each MAR system ideally needs a dedicated caretaker to keep it functional. We did not observe or interview the assigned caretakers but almost all MAR sites were functioning throughout the year. The caretakers assisted study field research assistants during water sample collection, pump motors were found operational, evidence of continued maintenance and continued community involvement which is promising for the future scale-up. Water quality monitoring may need further attention.

Notwithstanding these limitations, our results suggest that the MAR system significantly improved the microbiological quality of drinking water over pond water. Promoting MAR water consumption over pond water in salinity affected rural coastal communities potentially reduces the risk of waterborne disease.

## Acknowledgments

Financial support: The study has been funded by Wellcome Trust, UK (Grant no.106871/Z/15/Z), Our Planet, Our Health Award. Wellcome Trust reviewed and approved the design as the condition of providing funding but will not have a role in data collection, data analysis, data interpretation or writing of the report. The study was implemented by International Centre for Diarrhoeal Disease Research, Bangladesh (icddr,b). icddr,b acknowledges with gratitude the commitment of Wellcome Trust, UK, to its research efforts. icddr,b is also grateful to the Governments of Bangladesh, Canada, Sweden and the UK for providing core/unrestricted support. The icddr,b acknowledges with gratitude the study participants, the dedicated field team.

